# Variable E-field properties of dual-site tACS

**DOI:** 10.1101/2025.08.06.668900

**Authors:** Silvana Huertas-Penen, Maria Carla Piastra, Oula Puonti, Bettina C. Schwab

## Abstract

**Background:** Dual-site transcranial alternating current stimulation (ds-tACS) enables the modulation of interregional functional connectivity by introducing a phase lag between the stimulating currents. However, overlapping electric fields (E-fields), particularly in closely spaced cortical targets like the primary motor cortices (M1s), may unintentionally alter E-field characteristics and confound the interpretation of functional connectivity modulation.

**Objective:** We aimed to systematically evaluate how different phase lags affect key E-field characteristics when using high-definition ds-tACS, particularly when targeting the M1s. We sought to determine which montage configuration best preserved stable E-field characteristics and investigated whether individualised montage selection could enhance control over E-field consistency.

**Methods:** We used individualised finite-element method simulations based on MRI-derived head models to quantify the effects of different phase lags on E-field characteristics. E-field magnitude, normal component, spatial distribution, directionality, and effective stimulation area were assessed for nine montages and eight phase lags.

**Results:** All E-field properties, including E-field peak magnitude, peak magnitude of the normal component, redistribution, difference to optimal directionality, and effective area of stimulation, were modulated significantly across different phase lags for all tested montages. Furthermore, we found substantial inter-individual variability in all E-field properties. Individual selection of montages improved critical properties, particularly the E-field directionality.

**Conclusions:** In contrast to common assumptions, variations in the phase lag can significantly affect key E-field properties of high-definition ds-tACS. Therefore, we recommend considering modulations of the E-field characteristics when comparing physiological or behavioural effects of ds-tACS at different phase lags. Moreover, given the high inter-individual variability, we suggest the individualisation of montages to the most relevant E-field property.

**Highlights:**

- Dual-site tACS is often used to modulate functional connectivity.
- Changes in E-field characteristics with varying phase lags are undesirable.
- We used FEM models to quantify these changes across arbitrary phase lags.
- E-field characteristics of all montages varied significantly with the phase lag.
- Individualised montage selection was able to improve critical characteristics.

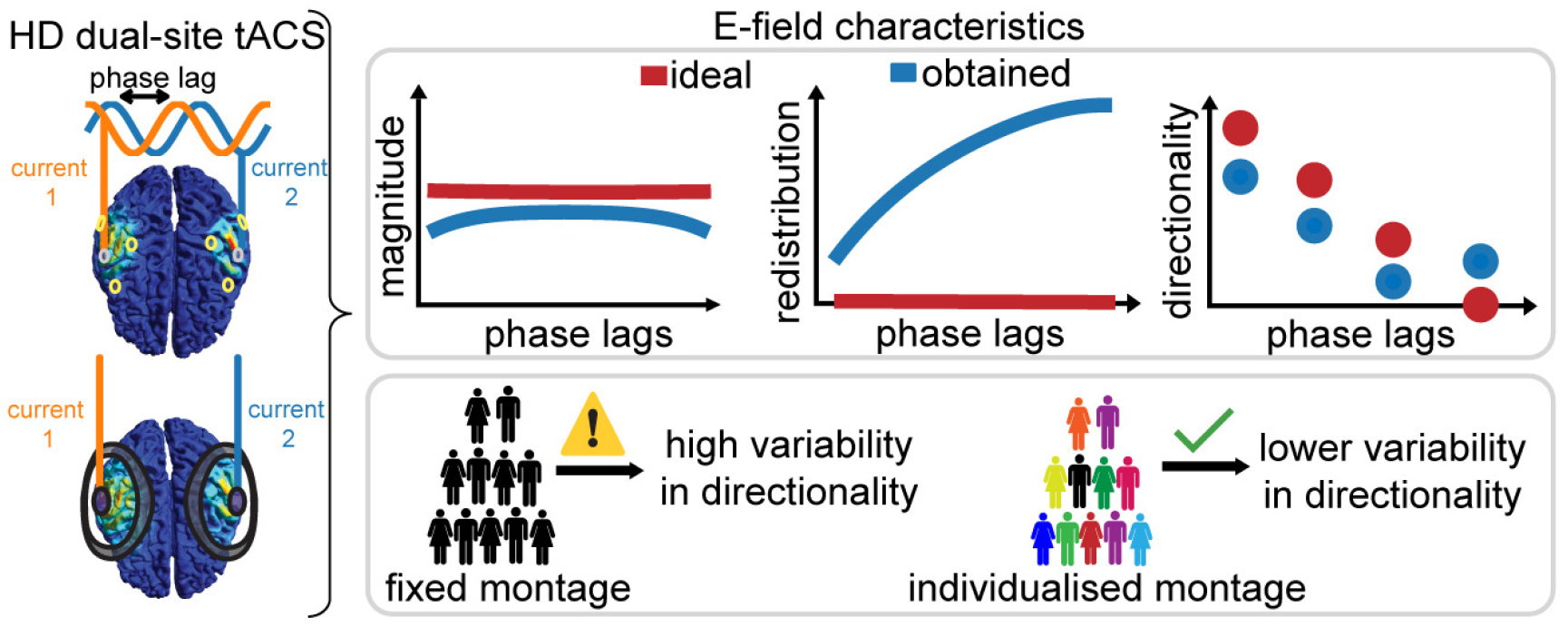

## 1. Introduction

Transcranial alternating current stimulation (tACS) noninvasively delivers alternating electric currents to the scalp, generating weak electric fields (E-fields) in cortical regions. These weak E-fields have recently been shown to phase-specifically modulate neural activity [1, 2, 3]. A specific variant, dual-site (ds) tACS, introduces a phase lag between the stimulation currents of the two targeted brain regions, changing the relative directionality of E-fields. This change in directionality enables the modulation of interregional functional connectivity (FC) [4, 5, 6, 7]. Such modulation is of great interest for both basic and clinical neuroscience, as it allows the investigation of the causal influences of FC on behaviour and may pave the way for readjusting pathological levels of FC in diseases. Ds-tACS has thus been widely applied, for example, for the modulation of FC between frontal and temporal regions [8] or motor cortical networks [9]. In sum, a number of studies have demonstrated that ds-tACS can lead to changes in FC and behaviour [8, 10, 11], even demonstrating robust correlations between the two [12, 13, 14, 15, 16, 17].

Nevertheless, ds-tACS is often applied with the assumption of spatially separated E-fields. Separated E-fields imply that phase lag manipulation would influence only the timing and, therefore, the relative directionality of the two E-fields, while leaving the E-field magnitude, spatial properties, and stimulated area unchanged. However, ds-tACS typically leads to at least partially overlapping E-fields. These overlaps can result in complex spatial interactions, where phase lag–dependent interactions unintentionally alter E-field characteristics [18]. Consequently, these changes in E-field properties may modulate not only FC but also other neural properties, such as oscillatory power, challenging the goal of selectively targeting FC.

To improve the spatial precision of stimulation and reduce unintended effects, high-definition (HD) montages have been developed [19]. Compared with conventional two-electrode setups, HD montages provide more focal stimulation [20, 21, 22]. While HD montages represent a significant advance, they do not fully eliminate the issue of overlapping E-fields [18, 21]. This overlap is particularly relevant when stimulating close-by brain regions such as the primary motor cortices (M1s). Although previous modeling work has described the problems caused by overlapping E-fields on HD montages [18], the quantitative extent to which phase lags affect E-field characteristics remains poorly understood. Furthermore, the E-field properties of phase lags beyond zero and *π* have not yet been investigated.

Thus, in this study, we aimed to systematically quantify the effects of different phase lags between stimulation currents at two sites on key E-field characteristics of HD ds-tACS. We focused on the M1s, which are common targets for neuromodulation [4, 9, 23, 24, 25, 26, 27, 28, 29, 30, 31]. We constructed individualised finite-element method (FEM) simulations based on MRI-derived head models using open-access data from the Human Connectome Project (HCP) [32]. We then assessed the phase lag-dependent variations in the E-field properties, including field magnitudes, spatial distribution, directionality, and effective stimulation area for nine montages and eight phase lags. Additionally, we sought to determine which HD montage configuration best preserved stable E-field characteristics, ensuring consistency in the stimulation effects. Finally, we investigated whether individualised montage selection could enhance control over E-field consistency and thereby improve the interpretability of ds-tACS interventions.

## 2. Material and Methods

### 2.1. Data set

We used data from the HCP [32], which included preprocessed T1 and T2 MRI scans at 3T [32]. 18 individuals (nine females) were randomly selected, with six from each age group: 22-25, 26-30, and 31-35 years. The IDs of the selected individuals are listed in Appendix A, Table A.1.

### 2.2. Electrode montages

We simulated nine typical motor cortex montages: five HD multiple small electrodes (MSE) montages (Figure 2B) and four HD ring electrode montages (Figure 2C). The ring montages varied in electrode size [5, 18, 33, 34], with the central electrodes positioned above the M1s (C4 in the right hemisphere and C3 in the left). Each ring montage included a ring electrode with varying inner and outer radii around the central electrode, as detailed in Appendix B, Table B.1. The electrodes were selected from commercially available sizes.

Two common MSE electrode configurations were used: a central electrode surrounded by three electrodes (3×1) [7, 8, 35, 36] or four electrodes (4×1) [6, 14, 18, 37, 38, 39, 40, 41, 42, 43]. The positions of the electrodes were based on the extended SPM12 EEG system 10-20, available in SimNIBS 4.0 [44, 45]. Table B.2 in Appendix B lists the electrode positions for each montage. All electrodes had a diameter of 12 mm and a height of 1 mm (StarStim, Neuroelectrics, Spain), except for those used in the 3×1c montage, which followed the specifications described by Gann et al. (2025) [7].

The central electrodes in both montage types delivered currents in the polarity opposite to those of the surrounding electrodes. In the MSE montages, the current of the central electrode was three or four times higher than that of the surrounding electrodes, whereas the currents were equal in the ring montages. This current distribution ensured a net current of zero. Table B.3 in Appendix B lists the conductivity values for each brain tissue layer [45] and for the electrodes used.

### 2.3. Simulation framework

Individualised head meshes (FEM models) were generated using the charm function of SimNIBS version 4.0 [45], which segmented the heads into nine compartments, and E-fields were simulated using SimNIBS [44]. Based on Saturnino et al.(2017) [18], the E-fields for the ds-tACS simulations were calculated using three tDCS simulations per dataset and montage based on linear superposition:

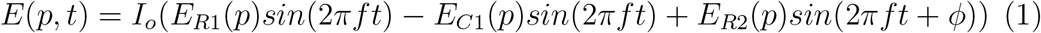

where *I_o_* is the amplitude of the applied current, *p* is the spatial position, *t* is the time, and *f* is the stimulation frequency. *E*_*R*1_(*p*) *E*_*C*1_(*p*), and *E*_*R*2_(*p*) denote the E-fields obtained from three separate tDCS simulations, each using a common return electrode (C2), located in the left hemisphere.

- *E*_*R*1_(*p*) corresponds to the E-field generated by currents between the return electrode (C2) and the surrounding right electrodes (R1) as shown in Fig. 1A.
- *E*_*C*1_(*p*) represents the E-field produced by current between the return electrode (C2) and the central right electrode (C1) as shown in Fig. 1B.
- *E*_*R*2_(*p*) corresponds to the E-field generated by currents between the return electrode and the surrounding left electrodes (R2) as shown in Fig. 1C.

**Figure 1.**
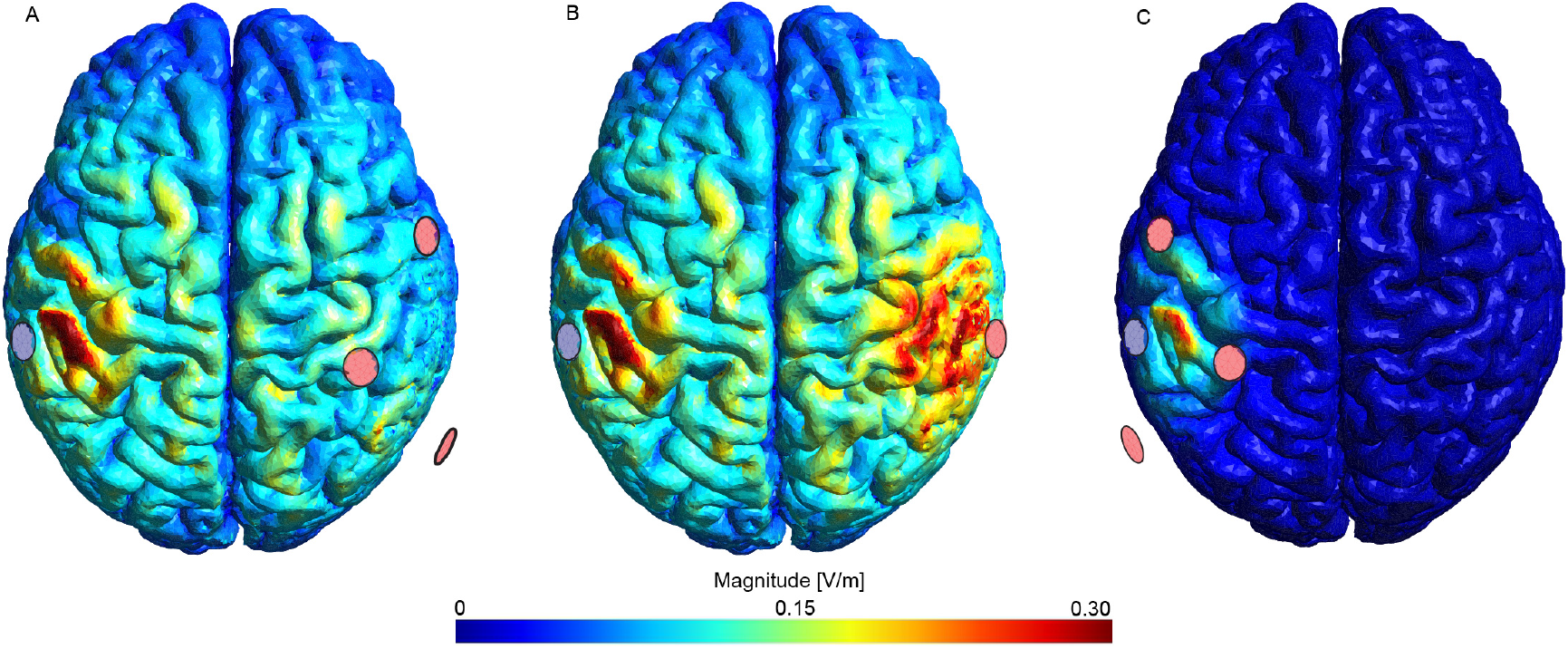
E-field magnitudes for the tDCS simulations with the central left electrode as the return common electrode, for an example individual (970764). A) *ER*1: Surrounding the electrodes in the right hemisphere. B) *EC*1: Central electrode in the right hemisphere. C) *ER*2: Surrounding electrodes in the left hemisphere. The red electrodes indicate the active sources (R1, C1, or R2), and the blue electrode represents the shared return electrode (C2). The E-fields of tACS were computed as linear superpositions of the depicted E-fields.

Phase lags were introduced by adjusting the phase of *E*_*R*2_, denoted as *ϕ*.

The E-fields were computed for 24 equally spaced time steps. This procedure is required when including phase lags other than zero and *π*, as in the remaining cases, there is not one time step where the maximum current is reached simultaneously in all channels. Thus, E-fields for all phase lags were simulated across all time steps, and the metrics were averaged across time steps for comparison (see Section 2.4).

The simulations were conducted with a maximum peak-to-peak current of 4mA and a frequency of 20 Hz. Because the E-fields scale linearly with the amplitude, our results can be directly translated into experimental settings with different current intensities. We explored eight phase lags ranging from 0 to 7/4*π*, thus equally sampling the entire cycle. Figure 2C illustrates sinusoidal stimulation for the first five phase lags; the remaining phase lags are symmetric around *π*. Figures 3A-D show the average E-field magnitude over time for one individual and two exemplary phase lags (0 and 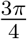) for one MSE montage and one ring montage.

**Figure 2.**
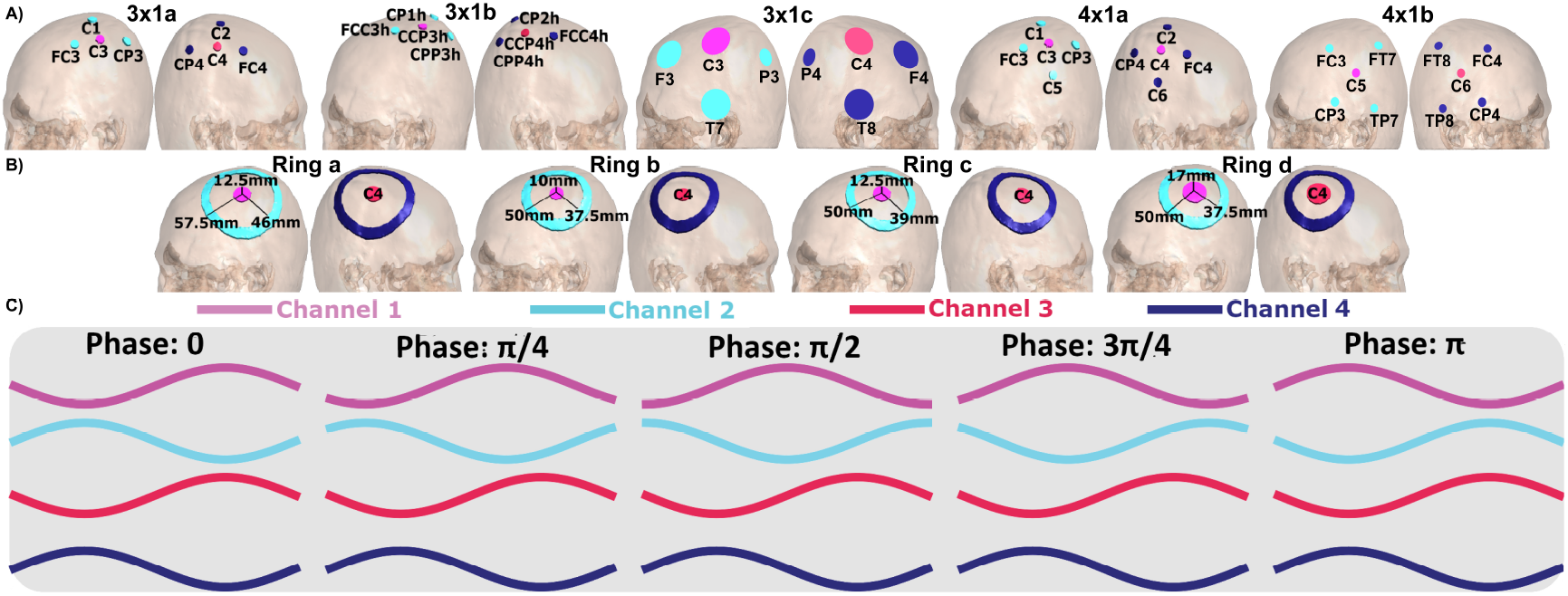
A) MSE montage configurations. B) Ring montage configurations. C) Injected currents with different phase lags.

**Figure 3.**
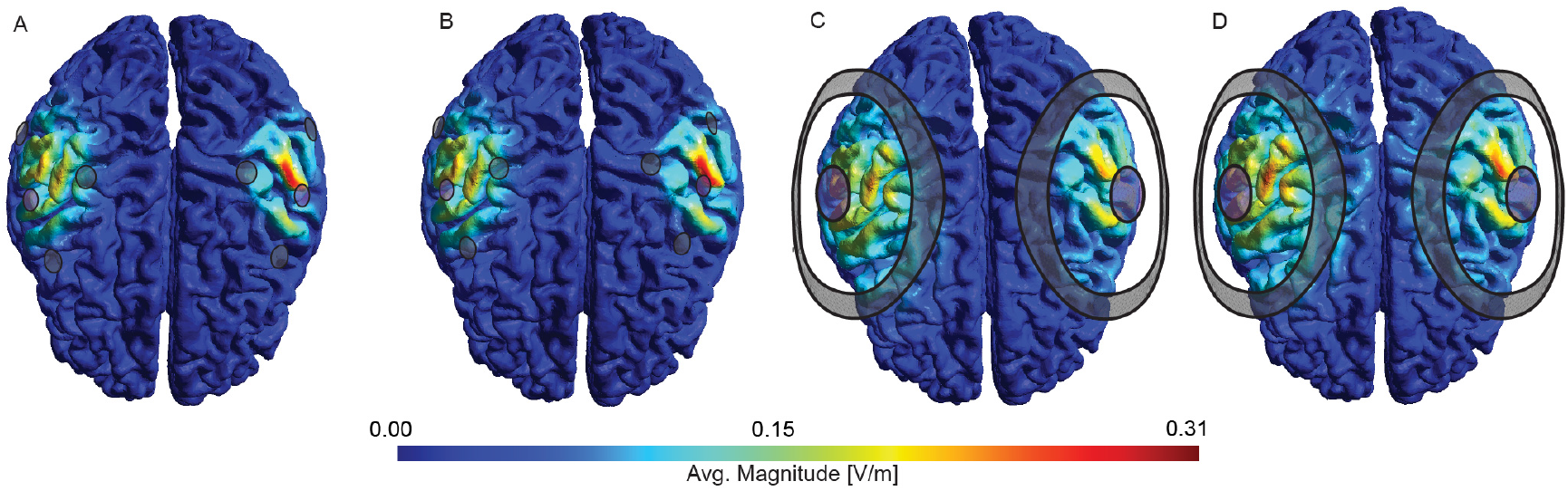
Time-average E-Field magnitudes for an example individual (970764). A) 3×1a MSE montage at phase lag zero B) and at phase lag 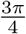. C) Ring b montage with a phase lag of zero; D) and a phase lag 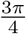. at one time step. The colour of the electrodes reflects current polarity

### 2.4. E-field metrics relevant to FC modulation

To quantify the E-field characteristics, we first removed outliers by excluding values above the 99.9th percentile of either the E-field magnitude or the absolute value of the component normal to the cortical surface (normal component), thereby minimising the influence of outliers [18, 46]. To further reduce the influence of numerical noise and potential inaccuracies in the SimNIBS simulations, we applied a lower threshold to both the E-field magnitude and its normal components. Values less than or equal to 2% of the absolute 99.9th percentile of the E-field were set to zero, suppressing negligible field contributions. The resulting data were used to compute the relevant metrics for each montage and phase lag.

For each individual, metrics were calculated for the grey matter sheet of the M1s and a surrounding gray matter sheet of 28 additional regions. The selected surrounding regions received an average E-field magnitude of 0.01 V/m or greater in all montages, both hemispheres, and all individuals (Fig. 4). Other regions were neglected in the analysis to ensure a physiologically meaningful field magnitude and to mitigate the impact of potential inaccuracies in the simulations. The regions were based on the Glasser Atlas [47]. A full list of the selected regions is provided in Appendix C, Table C.1.

**Figure 4.**
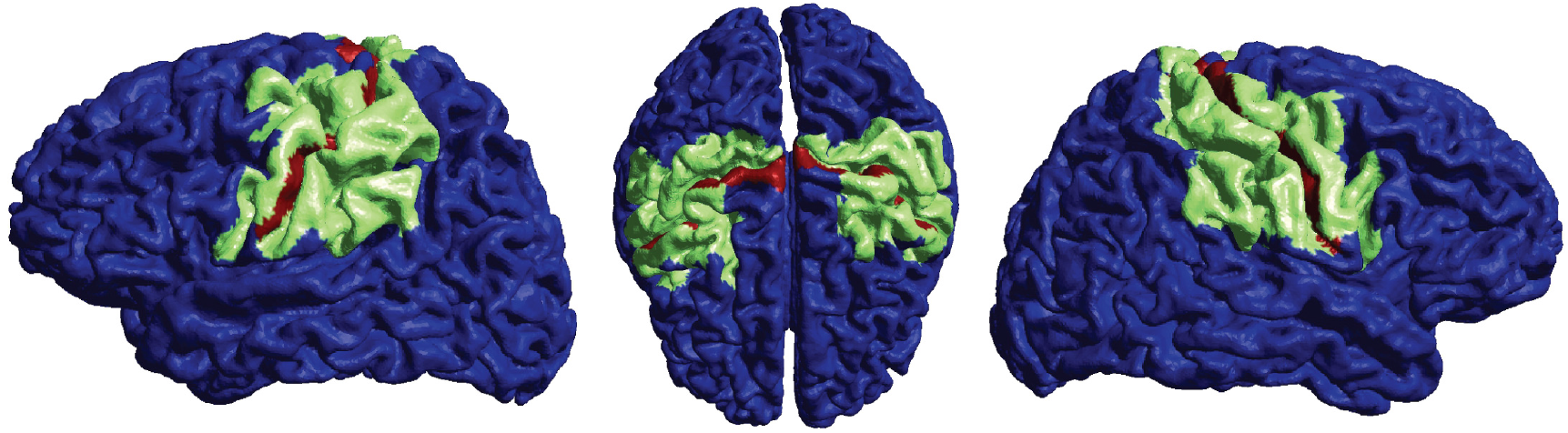
Surrounding gray matter sheet with minimum average E-field magnitudes of 0.01V/m (green), grey matter sheet of the M1s (red).

We investigated the following metrics:

#### A. Magnitudes

a. Peak magnitude of E-field: A commonly used measure for estimating the E-field strength [18].
b. Peak magnitude of E-field normal component: A commonly used measure for estimating the “effective” peak magnitude of the E-field, that is, in the direction of cortical dendrites, and thus normal to the grey matter sheet [18].

#### B. Redistribution

The relative distance measure (RDM) quantifies the dissimilarity between the spatial E-field distribution at a phase lag of zero and that at a nonzero phase lag [18, 48]:

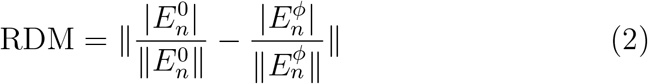

where 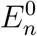 and 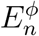 represent the E-field normal components of all nodes on the cortical surface for a phase lag of zero (0) and other phase lags (*ϕ*), respectively. The maximum value of the RDM is 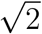, indicating that the normal component distributions of the two-phase lags do not overlap. An RDM value of zero signifies an identical distribution of the normal component of the E-field, which is the desired situation for ds-tACS.

#### C. Directionality

a. The Dot Product (DotP) quantifies changes in the directionality of the normal component of the E-field between the zero phase-lag condition and other conditions [18].

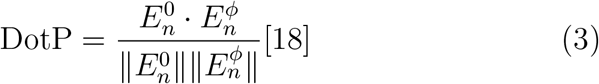 If the E-fields have the opposite directionality in all nodes, the Dot Product (DotP) is -1. By contrast, if the E-field has the same direction in all nodes, the DotP becomes 1.
b. Difference between the obtained and ideal DotP We define the difference between the ideal DotP (*IDotP*_0*,ϕ*_) and the obtained one (*ODotP*_0*,ϕ*_) between the E-fields of phase lags zero and *ϕ* as

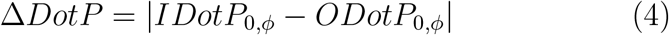

The ideal values were calculated as described in Appendix D.

#### D. Effective Area of Stimulation

We quantified the focality of the normal component of the E-field within the grey matter sheet of each M1 region using the effective area of stimulation [49]:

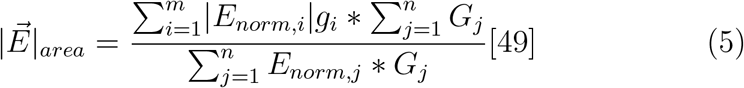

where *E_norm,i_* represents the normal components of the E-field that are equal to or greater than 50% of the peak normal component, and *g_i_* corresponds to the area of the nodes of these E-fields. *G_j_* denotes the area of all nodes in the region of interest, whereas *E_norm,j_* represents the normal components of all E-fields within the region of interest. Thus, the effective area of stimulation 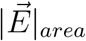 is the area in the target region where the normal components of the E-field are equal to or greater than 50% of the maximum E-field normal components on the surrounding grey matter sheet, including the M1s.

### 2.5. Variability of metrics

The coefficient of variation (CV) was used to quantify the inter-individual variability for each metric:

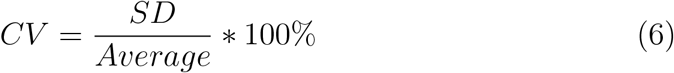

The standard deviation (SD) and average values across individuals were calculated for each metric and each montage, and then averaged across all phase lags.

Additionally, we tested the potential of individualising montages based on a preselected set of montages. The best-performing montage was selected for each individual. Both the metric of interest and its CV were then calculated across individuals using these selected montages and averaged over all phase lags. This test for individualisation was only performed for RDM and ΔDotP, the metrics which are specific and critical to dual-site tACS. In contrast, optimisation for magnitude and focality is also relevant for tACS in general and is already implemented in SimNINBS [38, 49, 50, 51].

### 2.6. Statistical Testing

For the peak E-field measurements, we selected the maximum value observed across all time steps. For the other metrics, we calculated the average value across time steps. To assess whether the metrics exhibited significant modulation across the phase lags, we used the weighted Hermans-Rasson (WHR) test [52]. The test was applied to each metric and montage for 10.000 iterations. To account for multiple montage comparisons, we applied the Bonferroni–Holm correction. A metric was considered significantly modulated by the phase lags if the corrected p-value was <0.05.

### 2.7. Spherical head model

In addition to the individualised head models, we simulated a spherical model, available in SimNIBS 4.0 [45], with a radius of 95 mm. This simplified model created a controlled environment for investigating the optimal E-field characteristics with minimal overlap of E-fields. Two montages were positioned at the maximum distance from each other on the sphere, each including a central circular electrode with a radius of 5 mm and a surrounding ring-shaped electrode with outer and inner radii of 15 and 10 mm, respectively. These montages were very small compared to the size of the sphere.

## 3. Results

### 3.1 Ideal values in a sphere model

We first investigated all metrics under ideal conditions with small ring electrode montages on a spherical model (Figure 5). As expected, the peak magnitude of the E-field normal components were low but stable over the phase lags. The RDM was almost zero for all the phase lags. Furthermore, the DotP was very close to the ideal value. In summary, under ideal conditions, it is possible to separate the two E-fields.

**Figure 5.**
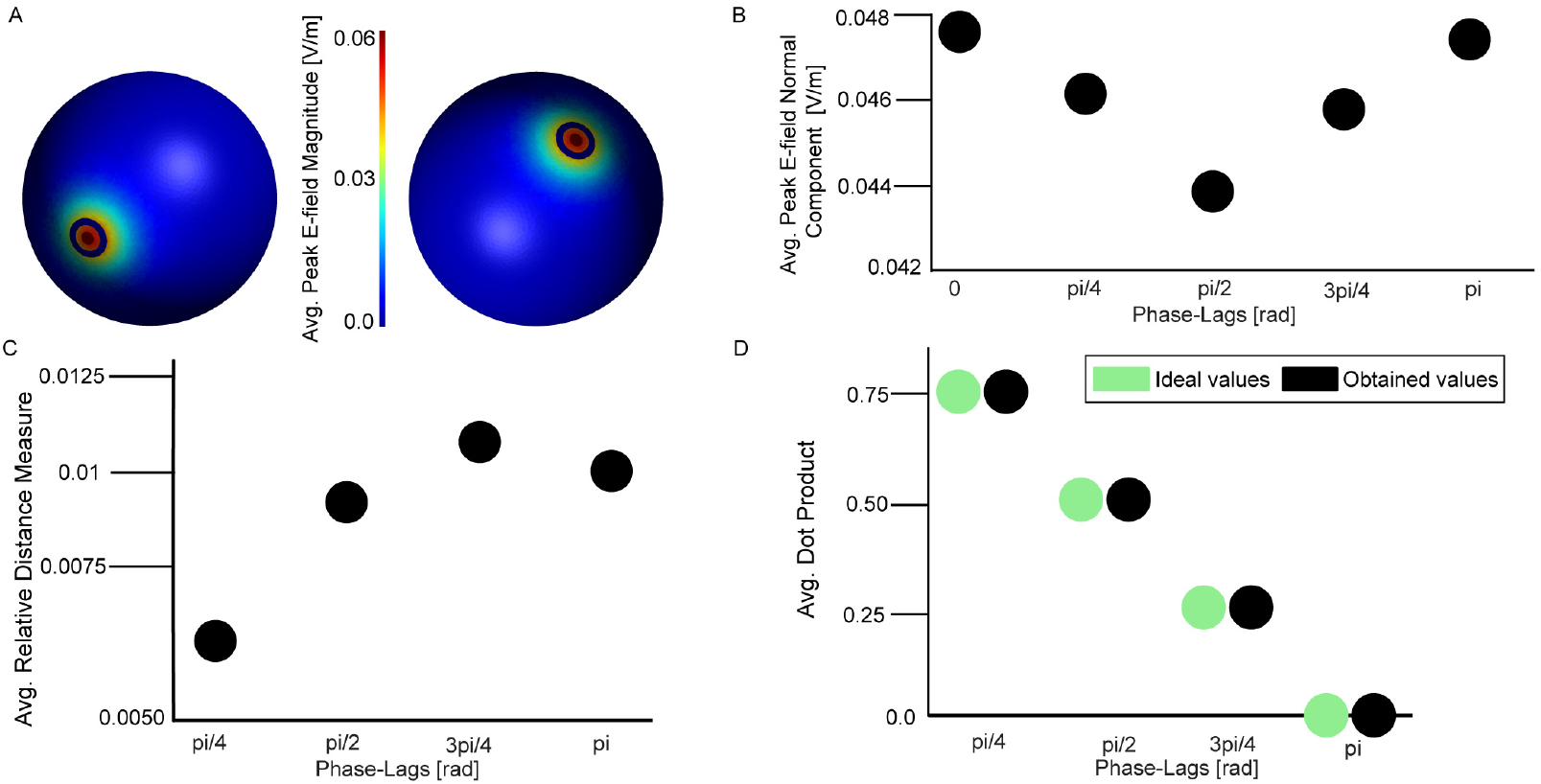
Near-ideal values for a sphere. A) Small ring electrodes were used on a large sphere, leading to low average E-field magnitudes. B) The average E-field magnitude normal component barely varies across the phase lags. C) The average RDM was low across all phase lags. D) The DotP (black) is close to the ideal (green) values for all phase lags.

### 3.2 E-field magnitudes slightly depend on the phase lag

Next, we studied the metrics for realistic head models. As the first key question, we examined how the E-field magnitude varied across the phase lags for different montages. Figures 6 and 7 show the peak E-field magnitude and peak inward normal component for all montages and individuals. Tables 3.1 and 3.2 present the average values per montage. Notably, different average properties, such as the largest mean and the smallest differences across phase lags, are optimal for different montages. In other words, no single montage maintained optimal E-field properties across all phase lags. The peak E-field magnitude and peak magnitude inward normal component, computed across all individuals and montages for both the M1s and the neighbouring regions, were slightly but significantly influenced by the phase lag (all corrected p-values < 10^−3^, WHR *>* 4.04 for the M1s and WHR *>* 0.51 for the surrounding regions).

**Table 3.1:** Overview of population-average peak E-field magnitudes per montage. The average/difference between the maximum and minimum across all phase lags and individuals are shown. The best montage for every measure (largest mean and smallest difference) is highlighted in green.

|  | 3x1a | 3x1b | 3x1c | 4x1a | 4x1b | Ring a | Ring b | Ring c | Ring d |
| --- | --- | --- | --- | --- | --- | --- | --- | --- | --- |
| Avg. Peak Magnitude Surrounding Regions [V/m] | 0.347 | 0.265 | 0.395 | 0.333 | 0.419 | 0.344 | 0.327 | 0.310 | 0.275 |
| Diff Max. - Min. Peak Magnitude Surrounding Regions [V/m] | 0.440 | 0.369 | 0.351 | 0.471 | 0.506 | 0.411 | 0.429 | 0.375 | 0.321 |
| Avg. Peak Magnitude M1s [V/m] | 0.278 | 0.221 | 0.360 | 0.252 | 0.332 | 0.319 | 0.298 | 0.284 | 0.257 |
| Diff Max. - Min. Peak Magnitude M1s [V/m] | 0.336 | 0.309 | 0.376 | 0.321 | 0.539 | 0.407 | 0.360 | 0.331 | 0.284 |

**Table 3.2:** Overview of population-average peak E-field normal components per montage. The average/difference between the maximum and minimum across all phase lags and individuals are shown. The best montage for every measure (largest mean and smallest difference) is highlighted in green.

|  | 3x1a | 3x1b | 3in1c | 4x1a | 4x1b | Ring a | Ring b | Ring c | Ring d |
| --- | --- | --- | --- | --- | --- | --- | --- | --- | --- |
| Avg. Peak Normal Surrounding Regions [V/m] | 0.260 | 0.192 | 0.310 | 0.252 | 0.328 | 0.270 | 0.255 | 0.243 | 0.215 |
| Diff Max. - Min. Peak Normal Surrounding Regions [V/m] | 0.346 | 0.280 | 0.289 | 0.362 | 0.422 | 0.329 | 0.341 | 0.300 | 0.263 |
| Avg. Peak Normal M1s [V/m] | 0.220 | 0.162 | 0.287 | 0.205 | 0.270 | 0.241 | 0.220 | 0.214 | 0.187 |
| Diff Max. - Min. Peak Normal M1s [V/m] | 0.243 | 0.256 | 0.292 | 0.226 | 0.404 | 0.312 | 0.257 | 0.252 | 0.214 |

**Table 3.3:** Overview of population-average RDM values per montage. The average/maximum across all phase lags are shown. The best montage for every measure (smallest value) is highlighted in green.

|  | 3x1a | 3x1b | 3x1c | 4x1a | 4x1b | Ring a | Ring b | Ring c | Ring d |
| --- | --- | --- | --- | --- | --- | --- | --- | --- | --- |
| Avg. RDM Surrounding Regions | 0.333 | 0.351 | 0.386 | 0.319 | 0.317 | 0.327 | 0.307 | 0.310 | 0.308 |
| Max. RDM Surrounding Regions | 0.396 | 0.416 | 0.497 | 0.388 | 0.413 | 0.408 | 0.383 | 0.389 | 0.386 |
| Avg. RDM M1s | 0.222 | 0.302 | 0.391 | 0.247 | 0.148 | 0.296 | 0.260 | 0.256 | 0.266 |
| Max. RDM M1s | 0.315 | 0.425 | 0.554 | 0.341 | 0.169 | 0.421 | 0.373 | 0.390 | 0.389 |

**Table 3.4:**
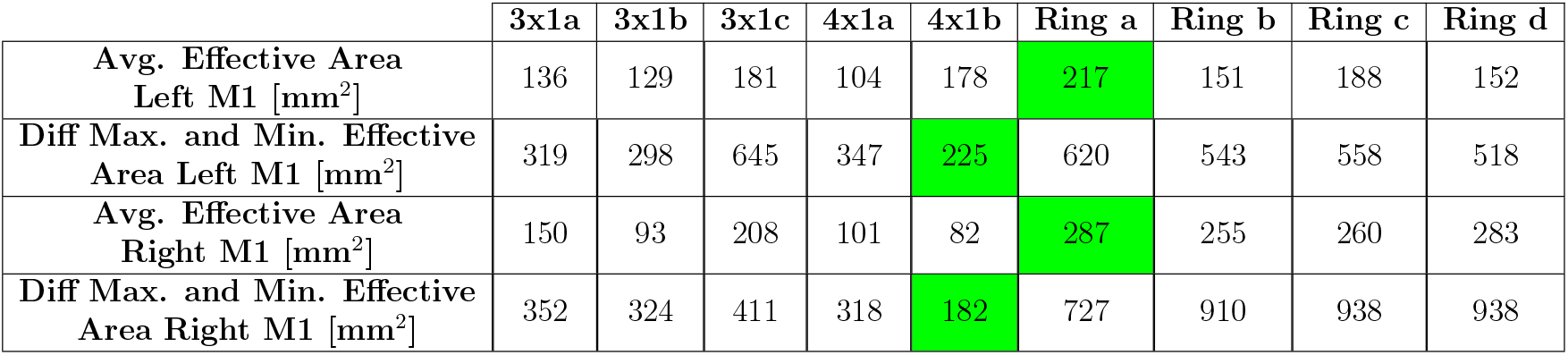
Overview of population-average effective area of stimulation per montage. The average/difference between maximum and minimum across all phase lags are shown. The best montage for every measure (largest mean and smallest difference) is highlighted in green.

**Table 3.5:**
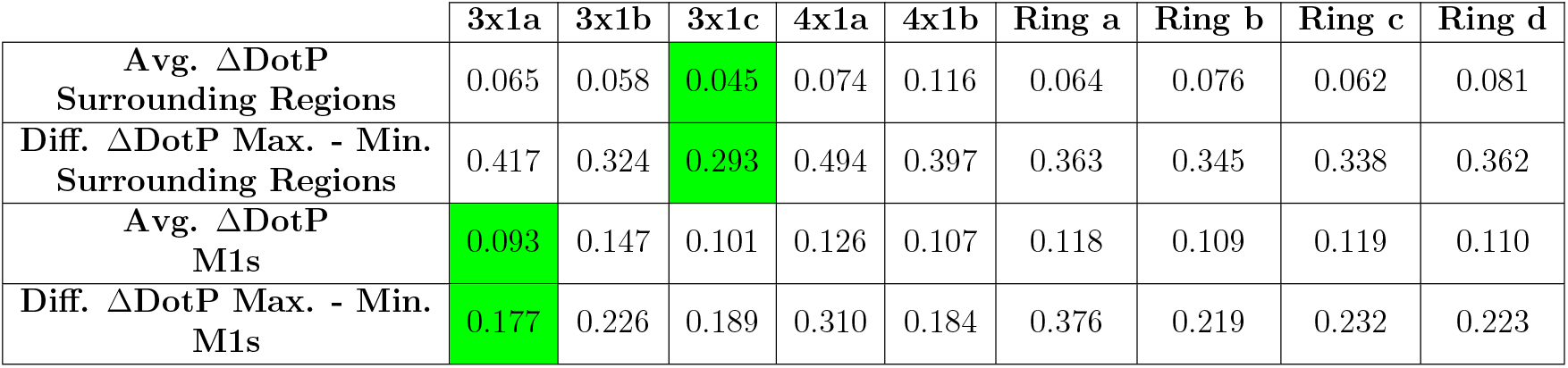
Overview of population-average ΔDotP. The average/difference between the maximum and minimum across all phase lags are shown. The best montage for every measure (smallest value) is highlighted in green.

|  | 3x1a | 3x1b | 3x1c | 4x1a | 4x1b | Ring a | Ring b | Ring c | Ring d |
| --- | --- | --- | --- | --- | --- | --- | --- | --- | --- |
| Avg. $\Delta\text{DotP}$ Surrounding Regions | 0.065 | 0.058 | 0.045 | 0.074 | 0.116 | 0.064 | 0.076 | 0.062 | 0.081 |
| Diff. $\Delta\text{DotP}$ Max. - Min. Surrounding Regions | 0.417 | 0.324 | 0.293 | 0.494 | 0.397 | 0.363 | 0.345 | 0.338 | 0.362 |
| Avg. $\Delta\text{DotP}$ M1s | 0.093 | 0.147 | 0.101 | 0.126 | 0.107 | 0.118 | 0.109 | 0.119 | 0.110 |
| Diff. $\Delta\text{DotP}$ Max. - Min. M1s | 0.177 | 0.226 | 0.189 | 0.310 | 0.184 | 0.376 | 0.219 | 0.232 | 0.223 |

**Table 3.6:** Overview of the CV for the different E-field characteristics, for each montage and for an average over all the montages. The montage with the lowest CV for each measure is highlighted in green.

|  | 3x1a | 3x1b | 3x1c | 4x1a | 4x1b | Ring a | Ring b | Ring c | Ring d | Avg. Montages |
| --- | --- | --- | --- | --- | --- | --- | --- | --- | --- | --- |
| CV Avg. Peak Magnitude Surrounding Regions [%] | 30 | 60 | 21 | 32 | 24 | 31 | 29 | 29 | 27 | 31 |
| CV Avg. Peak Magnitude M1s [%] | 29 | 32 | 21 | 31 | 31 | 30 | 28 | 28 | 27 | 29 |
| CV Avg. Peak Normal Surrounding Regions [%] | 29 | 29 | 20 | 31 | 23 | 31 | 28 | 29 | 27 | 28 |
| CV Avg. Peak Normal M1s [%] | 26 | 32 | 20 | 27 | 27 | 30 | 27 | 27 | 27 | 27 |
| CV Avg. RDM Surrounding Regions [%] | 19 | 16 | 15 | 22 | 26 | 20 | 25 | 25 | 24 | 21 |
| CV Avg. RDM M1s [%] | 20 | 19 | 21 | 22 | 26 | 18 | 18 | 19 | 19 | 20 |
| CV Avg. $\Delta\text{DotP}$ Surrounding Regions [%] | 106 | 92 | 96 | 112 | 63 | 102 | 106 | 103 | 104 | 98 |
| CV Avg. $\Delta\text{DotP}$ M1s [%] | 45 | 30 | 53 | 48 | 45 | 59 | 52 | 49 | 52 | 48 |
| CV Avg. Effective Area Left M1 [%] | 56 | 52 | 57 | 68 | 34 | 53 | 65 | 56 | 58 | 55 |
| CV Avg. Effective Area Right M1 [%] | 58 | 73 | 32 | 81 | 59 | 40 | 62 | 59 | 55 | 58 |

**Figure 6.**
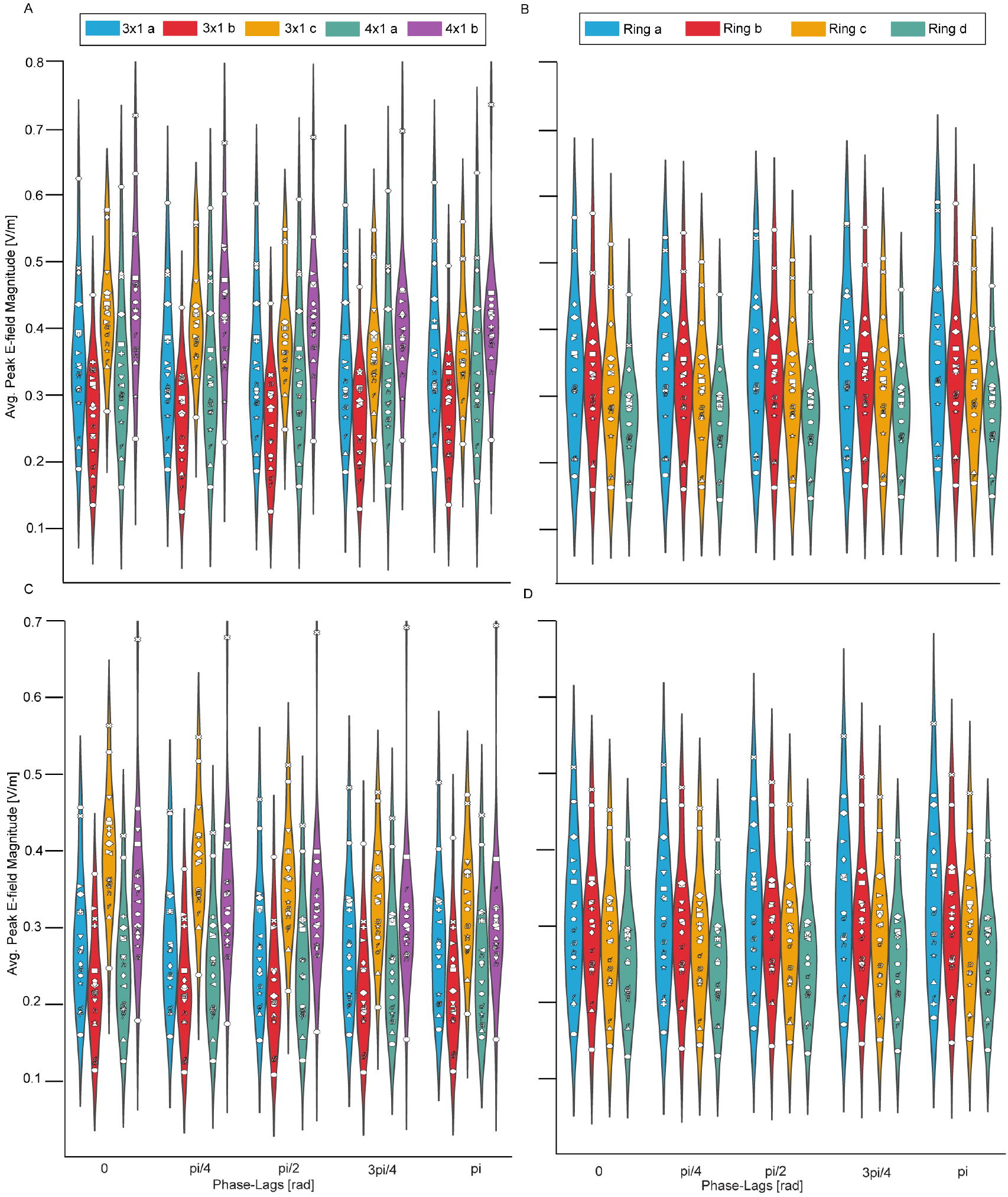
Maximum peak E-field magnitude for the MSE montages (A, C) and ring montages (B, D) in the surrounding grey matter sheet (A, B) and the M1s (C, D). The data from each individual are depicted as unique symbols.

**Figure 7.**
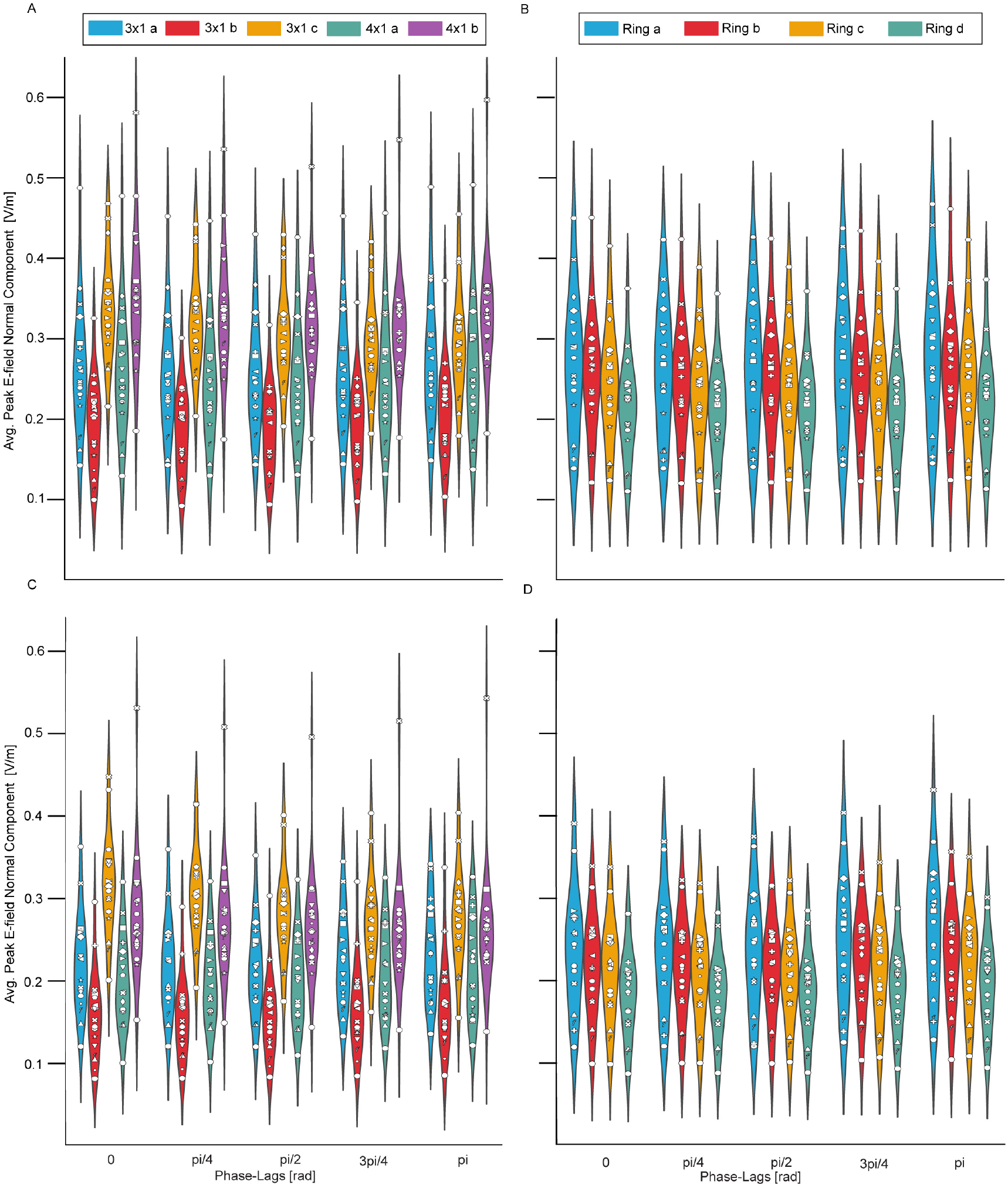
Maximum peak E-field normal components for the MSE montages (A, C) and ring montages (B, D) in the surrounding grey matter sheet (A, B) and the M1s (C, D). Data from every individual is depicted as a unique symbol.

### 3.3. E-field redistributions considerably change with the phase lag

The second key question focuses on investigating the extent to which the E-field distribution changes across phase lags. Figure 8 presents the RDM for the MSE and ring montages, comparing the dissimilarity in the E-field distribution between the zero-phase lag and the other phase lags. These comparisons were conducted across the surrounding grey matter sheet (Figures 8A-B) and grey matter sheet of the M1s (Figure 8C-D), with average properties described in Table 3.3. We observed a redistribution of E-fields from phase lag zero to all other phase lags, with the extent of this redistribution being strongly dependent on the phase lag for both the surrounding grey matter sheet and M1s (all corrected p-values < 10^−3^, WHR> 2.69 for the surrounding regions and WHR *>* 4.75 for the M1s).

**Figure 8.**
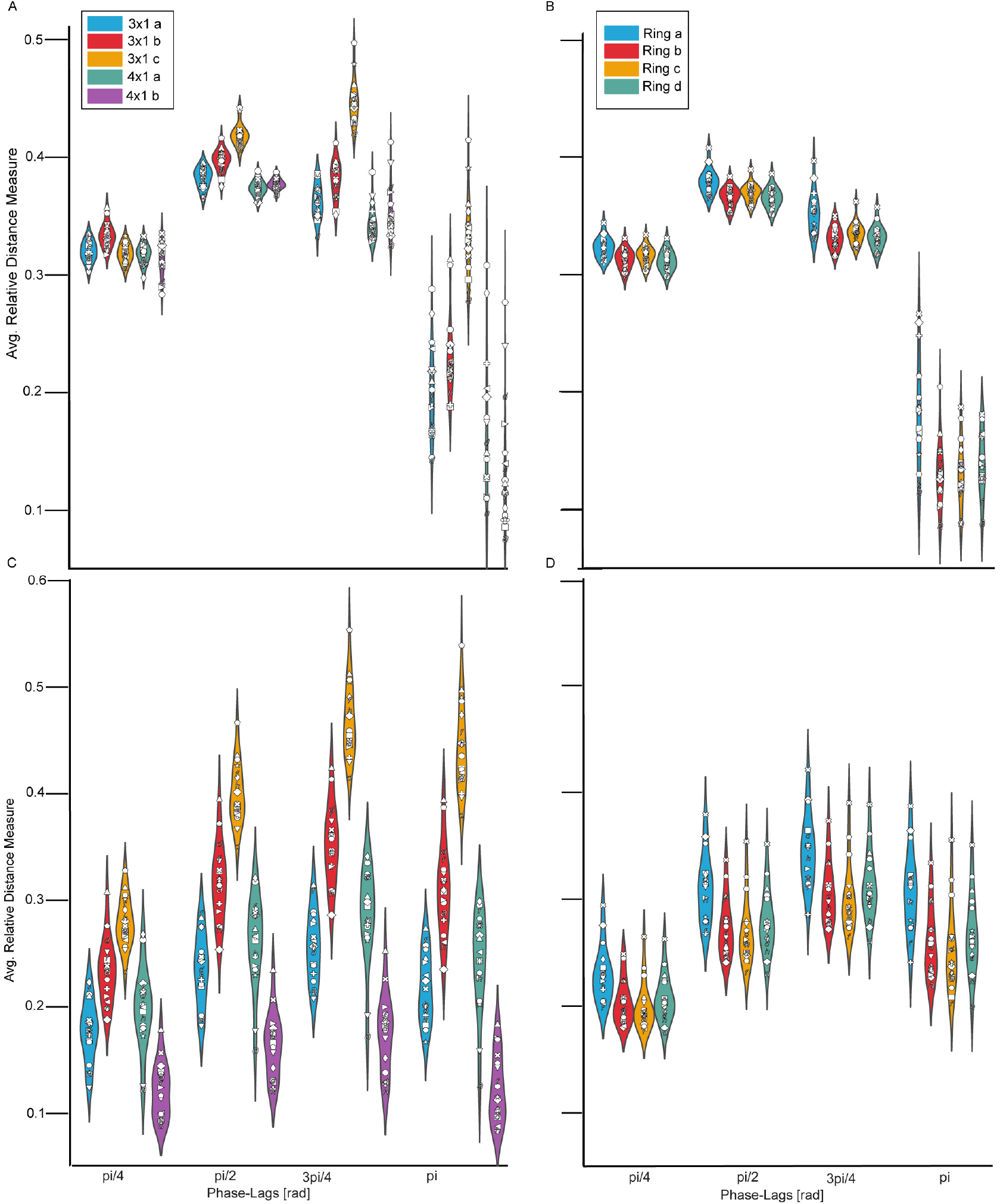
Average RDM for the the MSE montages (A, C) and ring montages (B, D) in the surrounding grey matter sheet (A, B) and in the M1s (C, D). The data from each individual are depicted as unique symbols.

### 3.4 The effective area of stimulation depends on the phase lag

As the third key question, we investigated the extent of tissue stimulated in the left and right M1s for each montage and how this metric varied across phase lags. Figure 9 illustrates the effective area of stimulation for the left (9A-B) and right(9C-D) M1s. The average metrics are listed in Table 3.4. For all montages, the effective area of stimulation significantly varied with the phase lag in the left hemisphere, right hemisphere, and average of both hemispheres (all corrected p-values< 10^−3^, WHR *>* 204729 for the left M1, WHR *>* 99750 for the right M1, and WHR*>* 1540601 for the average of both M1s).

**Figure 9.**
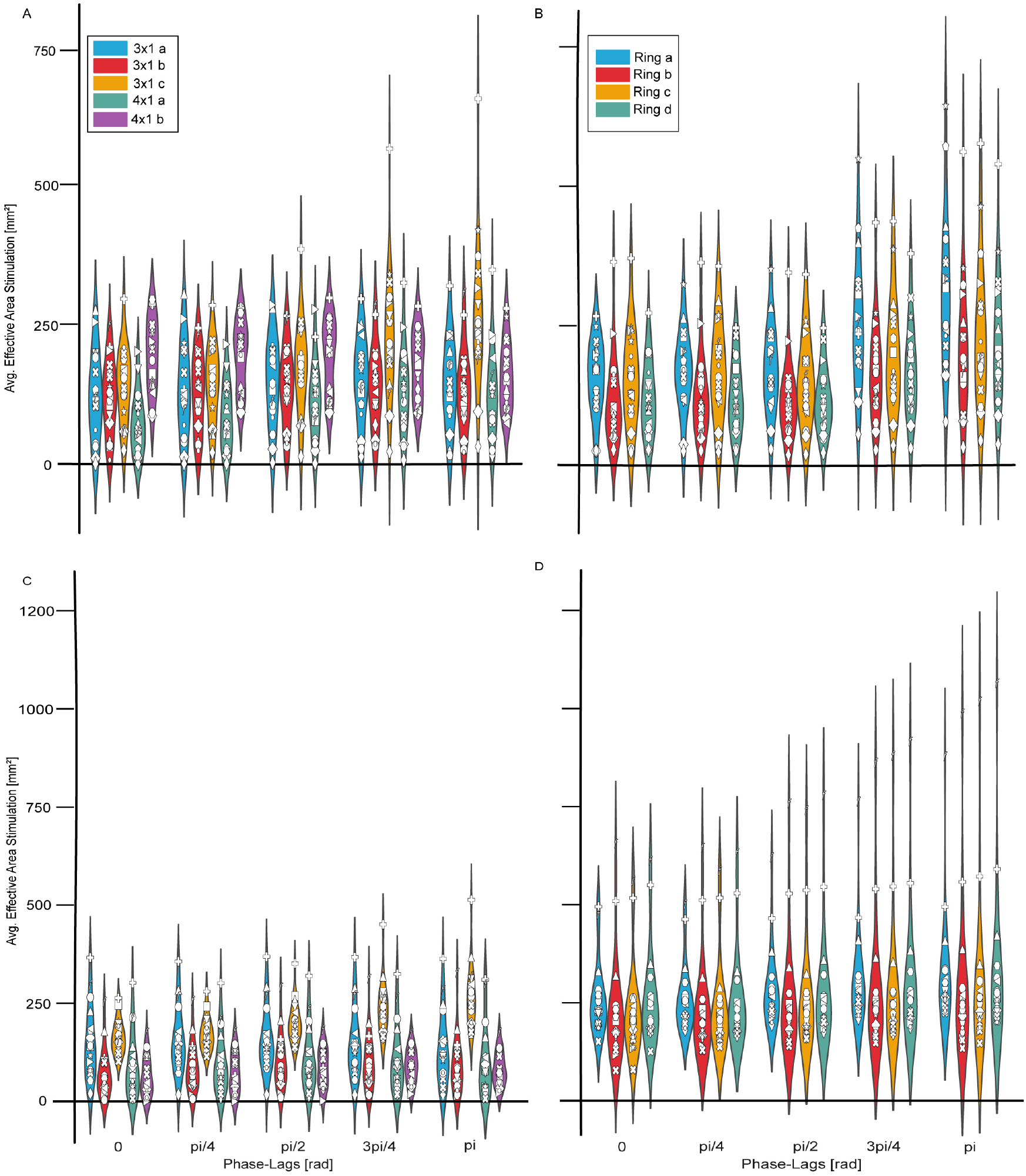
Average area of effective stimulation for the MSE montages in the left M1 (A) and right M1 (C), and for the ring montages in the left M1 (B) and right M1 (D). Data from every individual is depicted as a unique symbol.

### 3.5. Differences to the optimal E-field directions change with phase lags

The fourth key question examined the directionality of E-fields. Figure 10 illustrates the DotP between E-fields with a phase lag of zero and the other phase lags for both the surrounding grey matter sheet (10A-B) and the grey matter sheet of the M1s (10C-D) (see Table 3.5 for average measures). The ΔDotP showed a significant deviation from a uniform distribution for all montages (all corrected p-values < 10^−3^, WHR*>* 84.53 for the M1s, and WHR*>* 6.75 for the surrounding regions).

**Figure 10.**
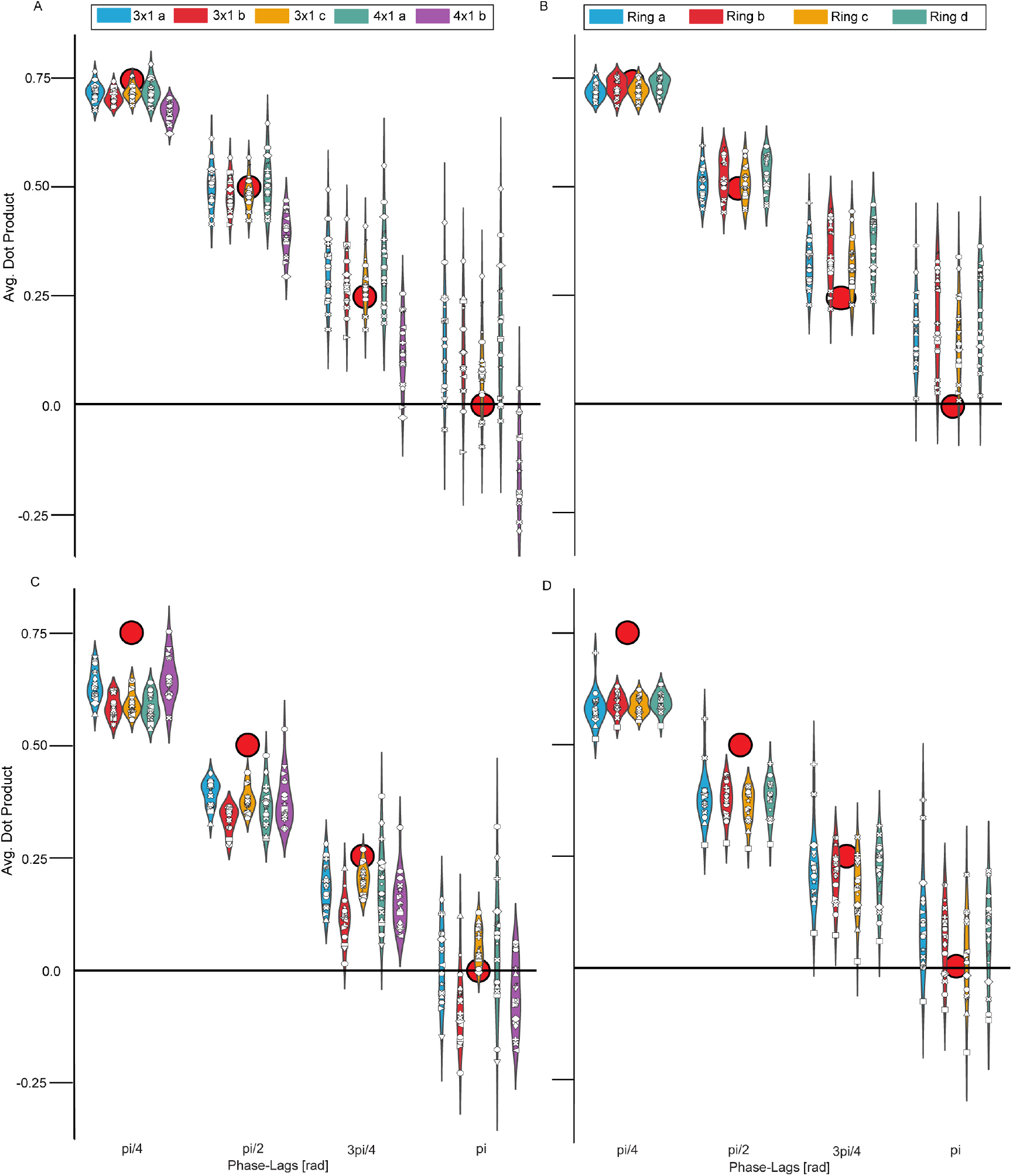
Average DotP for the MSE montages (A, C) and ring montages (B, D) in the surrounding grey matter sheet (A, B),and in the M1s (C, D). Data from every individual is depicted as a unique symbol. The red dots represent the ideal DotP values.

### 3.6. Substantial inter-individual variability

As the fifth key question, we quantified the variability of the investigated E-field properties across individuals. Table 3.6 lists the calculated CVs for all metrics. For all montages, the ΔDotP for the surrounding grey matter sheet exhibited the highest variability between the metrics, with an average CV, across all montages, of 98%. Also in the M1s, the average CV of the ΔDotP was 48%. Additionally, other metrics such as the effective area showed substantial inter-individual variability.

### 3.7 Potential for individualisation of montages

To evaluate the potential for individualising montages, we focused our analysis on the target area, the grey matter sheet of the M1s. For each metric, we first identified the montage that delivered the best results for the highest number of individuals, providing a group-level overview (see Supplementary Material 1). We then selected individual montages to assess an improvement in the overall RDM and ΔDotP as well as their variability.

At the group level, we compared all montages, including the MSE and ring montages, to determine which configuration was optimal for the E-field characteristics. The MSE 3×1c montage showed the greatest peak magnitude in 66.7% of the individuals. For the peak magnitude normal component, montages 3×1c and 4×1b showed the highest values for 44.4% of the individuals, each. Regarding the E-field redistribution, there was low variability, with 4×1b showing the lowest value for 94.4% of the individuals. Differences in the intended directionality were observed across individuals, with the 3×1b montage showing the lowest ΔDotP for 44.4% of the individuals. The effective area of stimulation showed high variability in the left M1, with 38.9% of individuals showing the highest value with Ring a. In contrast, 83.3% of the individuals showed the highest value in the right M1 with the same montage.

When considering only MSE montages, the MSE montage 3×1c produced the highest peak magnitude in the majority of individuals (66.7%). For the peak magnitude of the normal component of the E-field, montages 3×1c and 4×1b showed the highest values for 44.4% of individuals, each. The 4×1b montage also exhibited the most stable E-field redistribution (lowest RDM) for the largest fraction (94.4%) of the individuals. However, the montage with the lowest ΔDotP varied across individuals, with 3×1a showing the highest consistency for 44.4% of the individuals. Notably, montage 3×1c yielded the largest effective stimulated area for most individuals, for the left M1 (44.4%) and for the right M1 (72.2%).

For the ring montages, most individuals (94.44% and 88.8%) showed the highest peak and normal component magnitudes for Ring a, respectively. The lowest RDM values were observed for montage Ring c in 44.4% of the individuals. Additionally, the montage with the lowest ΔDotP in most individuals was the Ring d montage (44.4%). For the effective area of stimulation, Ring a showed the highest values in both M1s, with 83.3% of the individuals for the left M1 and 88.9% for the right M1.

To estimate the potential for individualisation of montages, we selected individual montages to minimise the ΔDotP or RDM for each individual and compared the results to the best-performing fixed montage. When optimising the ΔDotP, the average ΔDotP decreased from 0.15 to 0.08, and its CV decreased from 30% to 24%. When optimising the RDM, the average RDM decreased from 0.26 to 0.15, and its CV increased from 18% to 19%.

In summary, we found that certain fixed montages outperformed others, but no single montage was optimal for all metrics or all individuals. By selecting individual montages, the average values of RDM and ΔDotP decreased, indicating that the resulting E-field characteristics became more similar to the ideal case. Additionally, individualisation of montages reduced the variability of the ΔDotP.

## 4. Discussion

We systematically quantified the impact of varying phase lags on the E-field characteristics of ds-tACS, including its magnitude, redistribution, directionality, and effective area of stimulation. We observed significant differences in all key characteristics of the E-field across the phase lags and substantial inter-individual variability in all characteristics. When individualising montages to optimise the most critical characteristics, average directionality and average redistribution, these metrics could be improved by 40-50%.

Our findings indicate small but significant differences in the E-field magnitude and its normal component across different phase lags. Qualitatively, this finding has already been described by Saturnino et al. (2017) [18], who showed that for both a 4×1 and a ring montage, the E-field magnitude and focality varied between in-phase (zero) and anti-phase (*π*) conditions. Sim-ilarly, Alekseichuk et al. (2019) [53] experimentally confirmed the E-field magnitude differences between zero and *π* phase lags, although their study did not use an HD montage. We extend these findings to phase lags other than zero and *π*, and quantify the results for the case of targeting the M1s. Choosing arbitrary, for example physiologically plausible phase lags for ds-tACS may potentially boost the effects of ds-tACS [57, 7]. In the future, dynamic models may be used to estimate optimal phase lags for each individual [57, 58].

Even if small, potential differences in the E-field magnitude and other characteristics across phase lags are important when interpreting ds-tACS results. They may confound the phase-specific effects on FC by modulating other neural signals, such as EEG power or blood-oxygenation-level–dependent (BOLD) activity. Indeed, some ds-tACS studies reported phase-dependent modulation of EEG power. For example, Zhang et al. (2023) found both FC and power (ERD/ERS) differences between zero and *π* phase lags [54]. Similarly, Violante et al. (2017) and Preisig et al. (2021) reported phase-dependent changes in FC accompanied by BOLD activity modulation [12, 55]. In line with the relatively stable E-field magnitudes for HD montages observed in our study, most HD ds-tACS studies have reported changes in FC in the absence of power modulation [6, 7, 8, 56, 57]. It remains to be shown for every experimental study whether observed power differences may indeed arise from differences in E-field characteristics or could be functionally driven by differences in FC as an adaptive reaction.

As noted earlier [18], wider montages were associated with higher E-field magnitudes but also lower separability of E-fields (larger RDM and ΔDotP). This suggests a general trade-off between E-field magnitude and spatial specificity. Thus, for every experimental study, this trade-off should be considered. For example, clinical studies may profit from relevant effect sizes and may choose for larger E-fields at the cost of overlapping E-fields. On the other hand, basic science studies with many participants, which can deal with lower effect sizes, may rather profit from well-separated E-fields, avoiding confounding factors.

Next to changes across phase lags, also inter-individual variability is relevant for ds-tACS. While certain E-field properties, such as the RDM, were relatively stable across individuals, the ΔDotP and effective area of stimu-lation showed considerable inter-individual variability, next to the changes across phase lags. Thus, applying the same stimulation settings across individuals may result in inconsistent E-field directions and stimulated areas. In particular, the E-field directionality is crucial for connectivity modulation, and unintended deviations from the intended directionality could impede effective FC modulation.

We thus tested if individualisation of montages can also be done for E-field properties that are specific to ds-tACS, the redistribution of E-fields across phase lags, and the directionality of E-fields. Several studies already have demonstrated that differences in individual anatomy can lead to substantial variability in the magnitudes and focality of tACS, and that those features can be optimised using established pipelines [55, 59, 60, 61, 62]. In contrast, our approach for optimising the RDM and ΔDotP is based on the pre-selection of a number of montages which are all explicitly simulated, allowing to optimise for any arbitrary feature. We showed that individualising for the optimal directionality—a critical characteristic of ds-tACS, which is intended to drive the changes in FC—clearly improves the fit to the intended directionality, and reduces inter-individual variability. Such an individualisation may therefore enhance the precision and efficacy of ds-tACS.

Our study also shows some limitations. First, although our sample of 18 individuals was evenly distributed across age groups and balanced for biological sex, it may not fully capture the anatomical variability present in the general population. Similarly, our analysis was limited to healthy young individuals, and caution is warranted when generalising the findings to clinical or aging populations, where brain pathology may alter current flow. Nevertheless, we provide a methodology that can be used on any individual cohort and suggest a way to individualise montages. This is possible via the simulation of a range of montages, where the optimal montage can be chosen for each individual. Second, although our SimNIBS head models are detailed, they cannot capture dynamic effects, such as changes in skin conductivity due to sweating or movement of the brain within the CSF. Third, our optimisation approach, which is based on the explicit simulation of a number of pre-selected montages per individual, is computationally expensive. However, in the future, machine learning may help estimate individual E-fields without explicitly simulating them, strongly reducing the computational burden. Finally, our study described E-field characteristics, but cannot directly draw conclusions on the functional implications of these E-fields. Future studies may experimentally investigate whether the observed phase-lag-dependent differences in E-field characteristics translate to measurable differences in physiological or behavioural outcomes.

## 5. Conclusion

Our findings revealed that variations in the phase lag of HD ds-tACS significantly impact key E-field properties, introducing potential confounding factors for experimental designs. In particular, the relative directionality— the main factor intended to drive changes in FC—was different from the intended relative directionality and showed considerable inter-individual variability. The observed inconsistent E-field properties complicate the interpretation of physiological changes under ds-tACS.

In summary, we recommend considering phase-dependent changes in E-field characteristics like magnitude, distribution, directionality, and area of stimulation when designing ds-tACS studies, as they may potentially lead to unintended effects on neural activity. Using our methods, these properties can be simulated for any arbitrary phase lag, and can be statistically related to experimental outcomes. We furthermore propose that montages can be individualised based on a pre-selection of possible montages to optimise E-field characteristics for ds-tACS, and in particular, to achieve the desired directionality of E-fields.

## 6. Data availability statement

The code is available at https://gitlab.utwente.nl/bss_development/neuro/acs/EfieldPropertiesDualtACS. The raw data is available in the HCP database [32] under the S1200 database.

## 7. Acknowledgement

We would like to thank Axel Thielscher for very valuable discussions and advice, as well as Marieke Rona and Davide Liuni for help with the simulations.

## 9. Author roles

S. H. P.: Conceptualisation, Data curation, Formal analysis, Investigation, Methodology, Software, Validation, Visualisation, Writing — original draft. M. C. P.: Methodology, Writing — review and editing. O. P.: Formal analysis, Methodology, Software, Writing — review and editing. B. C. S.: Conceptualisation, Formal Analysis, Funding acquisition, Methodology, Project administration, Validation, Supervision, Writing — review and editing.

## Appendix A. IDs HCP datasets

**Table A1:**
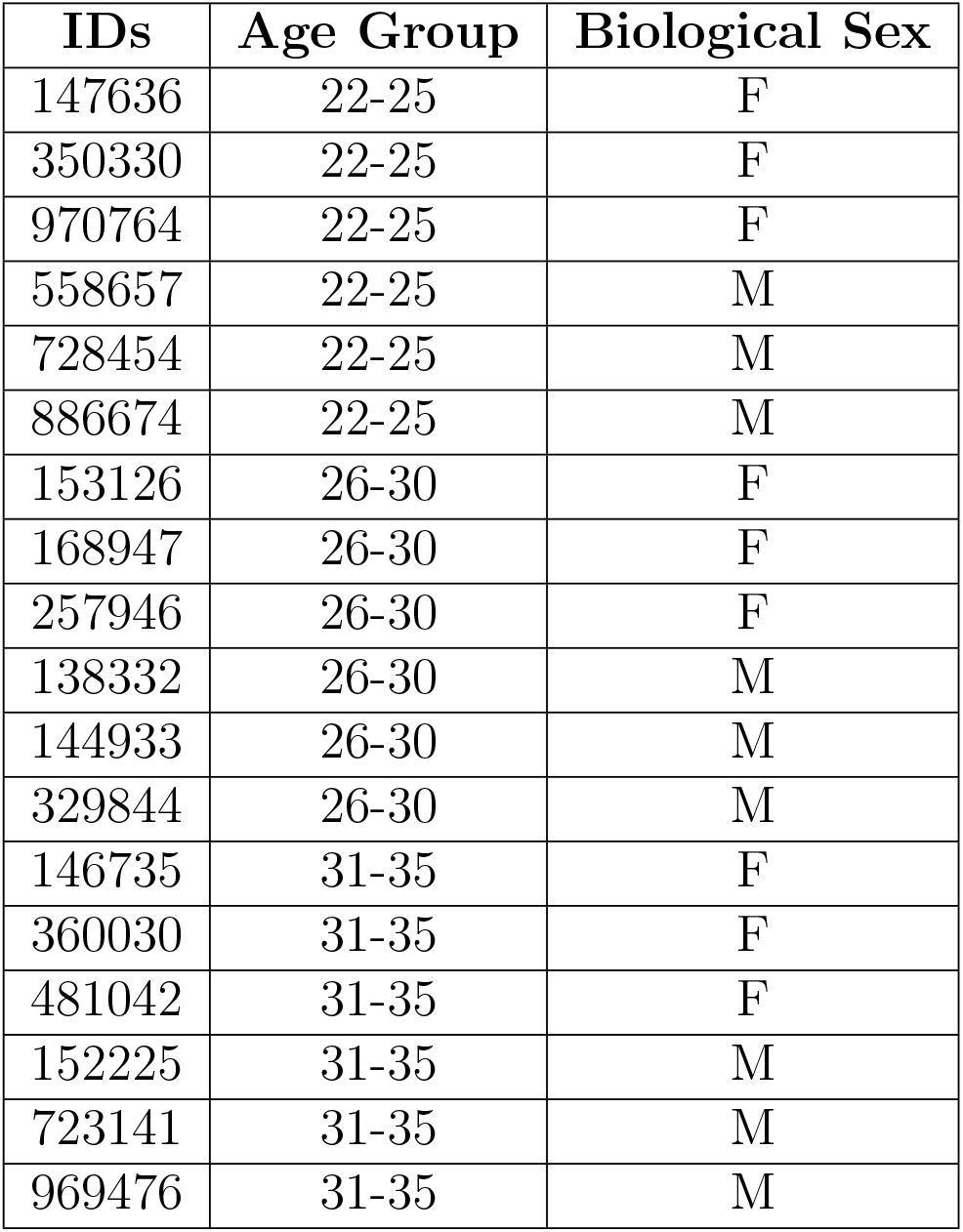
IDs of the datasets used for this study. The data was obtained from the HCP [32].

| IDs | Age Group | Biological Sex |
| --- | --- | --- |
| 147636 | 22-25 | F |
| 350330 | 22-25 | F |
| 970764 | 22-25 | F |
| 558657 | 22-25 | M |
| 728454 | 22-25 | M |
| 886674 | 22-25 | M |
| 153126 | 26-30 | F |
| 168947 | 26-30 | F |
| 257946 | 26-30 | F |
| 138332 | 26-30 | M |
| 144933 | 26-30 | M |
| 329844 | 26-30 | M |
| 146735 | 31-35 | F |
| 360030 | 31-35 | F |
| 481042 | 31-35 | F |
| 152225 | 31-35 | M |
| 723141 | 31-35 | M |
| 969476 | 31-35 | M |

## Appendix B. Montages and electrodes characteristics

**Table B.1:** Sizes of the ring montages used in the study. The table provides the radius of the central electrode and the inner and outer radii of the surrounding electrodes for different ring montage configurations.

| Ring Montages | Radius Central Electrode (mm) | Radii Surrounding Electrode (mm) |
| --- | --- | --- |
| Ring a | 12.5 | Inner: 46, Outer: 57.5 |
| Ring b | 10 | Inner: 37.5, Outer: 50 |
| Ring c | 12.5 | Inner: 39, Outer: 50 |
| Ring d | 17 | Inner: 37.5, Outer: 50 |

**Table B.2:** Positions of electrodes for the MSE montages. The table lists the central and surrounding electrode positions for different montage configurations, using the SPM12 EEG system.

| MSE Montages | Position Central Electrodes | Position Surrounding Electrodes |
| --- | --- | --- |
| 3in1a | C3, C4 | LH: C1, FC3, CP3. RH: C2, FC4, CP4 |
| 3in1b | CCP3h, CCP4h | LH: CCP1h, FCC3h, CPP3h. RH: CCP2h, FCC4h, CPP4h |
| 3in1c | C3,C4 | LH:F3, T7, P3 . RH: F4, T8, P4 |
| 4in1a | C3, C4 | LH: C1, FC3, C5, CP3. RH: C2, FC4, C6, CP4 |
| 4in1b | C6, C5 | LH: FT7, FC3, CP3,TP7. RH: FT8, FC4, CP4,TP8 |

**Table B.3:** Material and electrical characteristics of the electrodes and gel layers used in the study. The table lists the material composition, thickness, and conductivity values of both the MSE (Ag/AgCl) and the ring electrodes (rubber). The gel layer properties, including thickness and conductivity, are also provided.

| Parameter | Values |
| --- | --- |
| Thickness Electrodes | 1 mm |
| Thickness Gel Layer | 2 mm |
| Conductivity Gel Layer | 1 S/m |
| Conductivity Ring Electrodes and 3x1b (Silicone Rubber) | 29.4 S/m |
| Conductivity MSE (Ag/AgCl) except 3x1b | 13.6 S/m |

## Appendix C. Analysed regions

Table C.1 presents the 14 unilateral regions used for the analysis, all of which were derived from the Glasser Atlas [47]. The analysis included both hemispheres, resulting in 28 brain regions.

**Table C1:** Regions of the “surrounding grey matter sheet”. All regions were included bilaterally.

| Name Regions | Area Description |
| --- | --- |
| 1 | Area 1 |
| 2 | Area 2 |
| 3a | Area 3a |
| 3b | Primary Sensory Cortex |
| 55b | Area 55b |
| 6a | Area 6 Anterior |
| 6d | Dorsal Area 6 |
| 6r | Rostral Area 6 |
| 6v | Ventral Area 6 |
| FEF | Frontal Eye Fields |
| OP1 | Area OP1/SII |
| PF | Area PF Complex |
| PFop | Area PF Opercular |
| Pft | Area Pft |

## Appendix D. Calculation of the Ideal DotP values

The ideal DotP is determined under the assumption that two distinct areas are stimulated without leakage or overlap of the E-fields between the electrodes. Under these conditions, the E-fields in both areas were considered to be independent.

For ds-tACS, a phase shift is applied to the current in one area (*Area_a_*), whereas no phase shift was applied to the current in the other area (*Area_b_*). The applied currents are given by

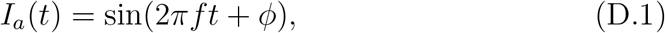

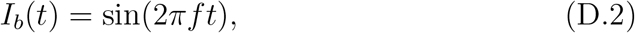

When calculating the DotP, we compared the direction of the normal component of the E-field at a given phase lag with that of the E-field at zero phase lag. In an ideal case, the direction of the normal E-field component is directly related to the applied current. Specifically, a positive current generates a positive normal E-field component (oriented inward towards the grey matter), whereas a negative current produces a negative normal E-field component (oriented outwards from the grey matter) [63].

In all cases, *Area_a_* did not receive a phase shift. Therefore, the normal E-field component in this area remained unchanged between the zero-phase and phase-lagged models. Consequently, the DotP for this area is always 1. In contrast, in *Area_b_*, where a phase shift is applied, the direction of the normal E-field component varies over time as a function of the phase shift *ϕ*:

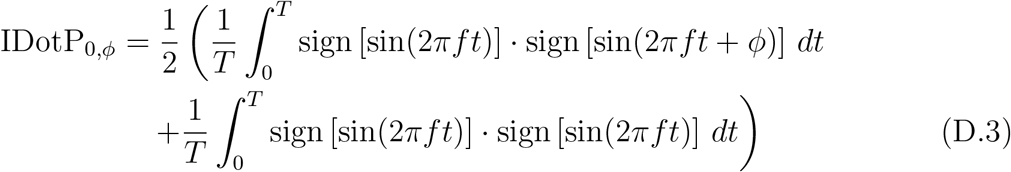

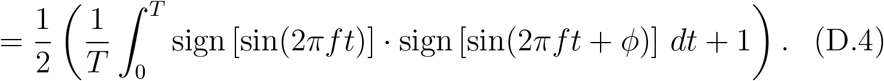

Here, T denotes the period of the sinusoidal function, and the division by two accounts for averaging across the two hemispheres or brain areas analysed. Importantly, the phase shift *ϕ* is applied only in one area.

To illustrate this, we calculated the ideal DotP for the *π/*2 phase lag condition. In *Area_a_*, the current remained identical in both the zero-and *π/*2-phase lag models, resulting in a constant direction of the normal E-field component throughout the sinusoidal cycle (Figure D.11A). Therefore, the DotP for this region is 1 (*IDotP_a_*). In contrast, in *Area_b_*, the sinusoidal currents for the zero-phase lag sin(2*πft*) and sin(2*πft* + *π/*2) differ in phase. Owing to this phase shift, the direction of the normal E-field component in *Area_b_* changes over time. In the ideal case, at certain time points, both currents share the same sign (either positive or negative; see Figure D.11B), implying that the normal E-field components are aligned, yielding an ideal DotP value of +1. At other times, the currents have opposite signs (one positive and one negative), indicating anti-alignment and resulting in an ideal DotP value of *−*1. At four specific time points, the current was zero, resulting in an ideal DotP value of 0.

**Figure D.11:**
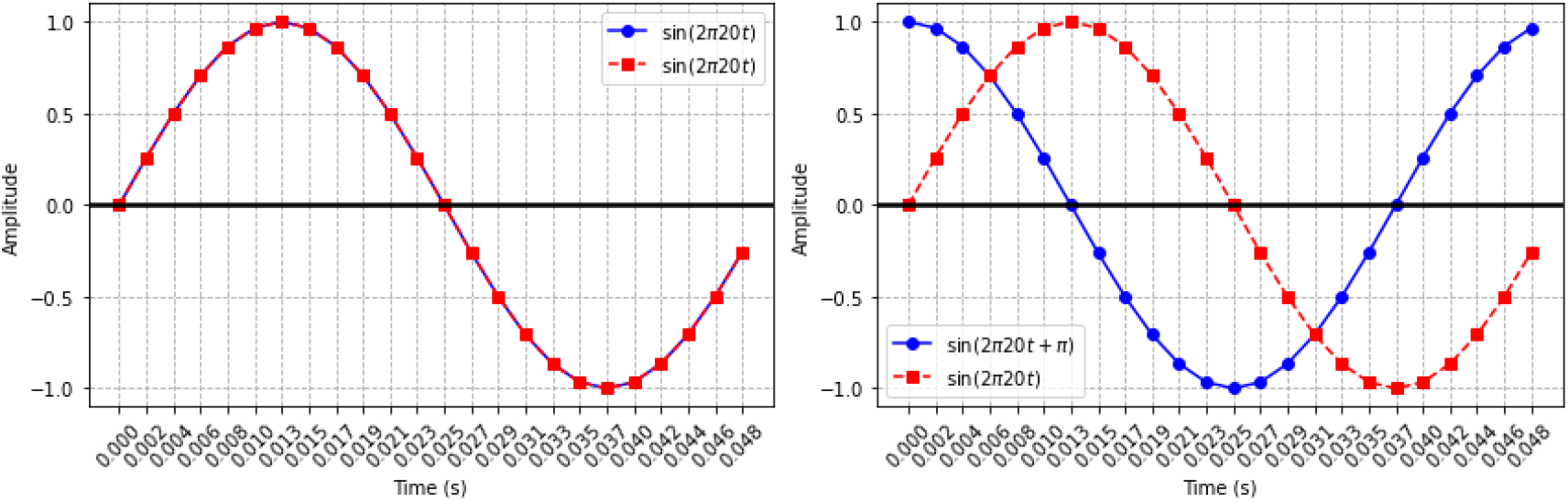
Example for computing the DotP under ideal conditions and a phase lag of *π/*2. A: Currents for *Areaa*. B: Currents for *Area_b_*.

In our simulations, we discretised the sinusoidal wave into 24 time steps. The ideal DotP was computed as follows:

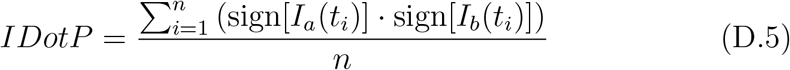

where:

- *I_a_*(*t_i_*) and *I*_2_(*t_b_*) represent the current at time step *t_i_* for the two regions,
- sign(*x*) denotes the sign of *x*, i.e., sign(*x*) = +1 if *x >* 0, sign(*x*) = *−*1 if *x <* 0, and sign(*x*) = 0 if *x* = 0,
- *n* is the total number of time steps.

For the discretised sinusoidal example, the IDotP in *Area_b_* becomes

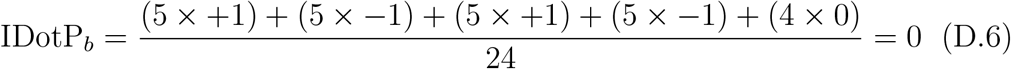

To obtain the ideal DotP, we averaged the ideal DotP values from both areas as follows:

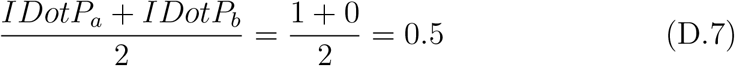

where *IDotP_a_* is the ideal DotP in one area and *IDotP_b_* is the ideal DotP in the other area. Thus, the ideal DotP for the *π/*2 phase lag condition is 0.5.

